# Disentangling shared and group-specific variations in single-cell transcriptomics data with multiGroupVI

**DOI:** 10.1101/2022.12.13.520349

**Authors:** Ethan Weinberger, Romain Lopez, Jan-Christian Hütter, Aviv Regev

## Abstract

Single-cell RNA sequencing (scRNA-seq) technologies have enabled a greater understanding of previously unexplored biological diversity. Based on the design of such experiments, individual cells from scRNA-seq datasets can often be attributed to non-overlapping “groups”. For example, these group labels may denote the cell’s tissue or cell line of origin. In this setting, one important problem consists in discerning patterns in the data that are shared across groups versus those that are group-specific. However, existing methods for this type of analysis are mainly limited to (generalized) linear latent variable models. Here we introduce multiGroupVI, a deep generative model for analyzing grouped scRNA-seq datasets that decomposes the data into shared and group-specific factors of variation. We first validate our approach on a simulated dataset, on which we significantly outperform state-of-the-art methods. We then apply it to explore regional differences in an scRNA-seq dataset sampled from multiple regions of the mouse small intestine. We implemented multiGroupVI using the scvi-tools library [1], and released it as open-source software at https://github.com/Genentech/multiGroupVI.

## 1 Introduction

The emergence of single-cell RNA sequencing (scRNA-seq) technologies has enabled the study of cellular heterogeneity at an unprecedented resolution [2]. To facilitate the investigation of various biological hypotheses, single-cell data are often collected simultaneously from multiple disjoint “groups”. For instance, recent studies have profiled epithelial cells from multiple regions of the mouse small intestine [3], cancer cell lines from multiple cancer types [4], as well as the response of innate immune cells to stimuli across multiple species [5]. In such scenarios, an important task is to isolate factors of variation that are shared across groups versus those that are specific to a given group. In the example of [4], Kinker et al. studied the transcriptomic heterogeneity within diverse cancer cell lines and their relationships to tumors, with the goal of further understanding drug resistance as well as metastasis in cancer.

However, few existing works address this problem. The analysis from the authors in [4] is ad-hoc, and consisted in performing non-negative matrix factorization separately on each cell line and then inspecting the learned factors to identify similarities as well as differences across cell lines. This pipeline involves several manual steps, which further highlights the need for a dedicated approach. Two previously proposed methods, multi-study factor analysis (MSFA) [6] and generalizable matrix decomposition framework (GMDF) [7] explicitly separate shared and group-specific variations in gene expression data. However, both are based on linear latent variable models and therefore have limited modeling flexibility compared to deep generative models (DGMs) [8; 9]. Davison et al. [10] recently proposed the cross-population variational autoencoder (CPVAE), a DGM for multi-group analysis, with a focus on image data. Yet, CPVAE makes a strong assumption about the data generating process by constraining the decoder network to be the sum of a nonlinear function of shared latent variables and a separate nonlinear function of group-specific latent variables. Such constraints may typically not hold in practice. Moreover, CPVAE as originally proposed does not account for the specific biases and noise characteristics of scRNA-seq data, a crucial condition for learning meaningful representations [11; 12].

A closely related line of work is concerned with so-called contrastive latent variable models [13; 14; 15; 16] that isolate factors of variation shared between a case and control scRNA-seq dataset from those that are enriched only in the case data. Previous work [13; 14] has demonstrated that isolating these case-specific factors of variation can reveal patterns in case where cells’ gene expression are difficult to discern using standard scRNA-seq analysis workflows. However, contrastive models require an explicit case-control relationship between datasets, which is not compatible with typical multi-group analysis scenarios (e.g. analyzing data from multiple tissues).

To address this gap in methodology, we developed multi-Group Variational Inference (multiGroupVI), a DGM that explicitly decomposes the gene expression patterns in scRNA-seq data into shared and group-specific factors of variation. For a scRNA-seq dataset with Γ disjoint groups, multiGroupVI models the variations underlying the data using Γ + 1 sets of latent variables: the first set, denoted as the *shared* variables, captures patterns shared across all groups while the remaining Γ sets model variations specific to individual groups. A graphical representation of the model in the case of two groups appears in **Figure 1**. To encourage the different sets of latent variables to exclusively capture their corresponding variations of interest, we make use of parameter sharing as well as a novel regularization term based on optimal transport [17].

**Figure 1:**
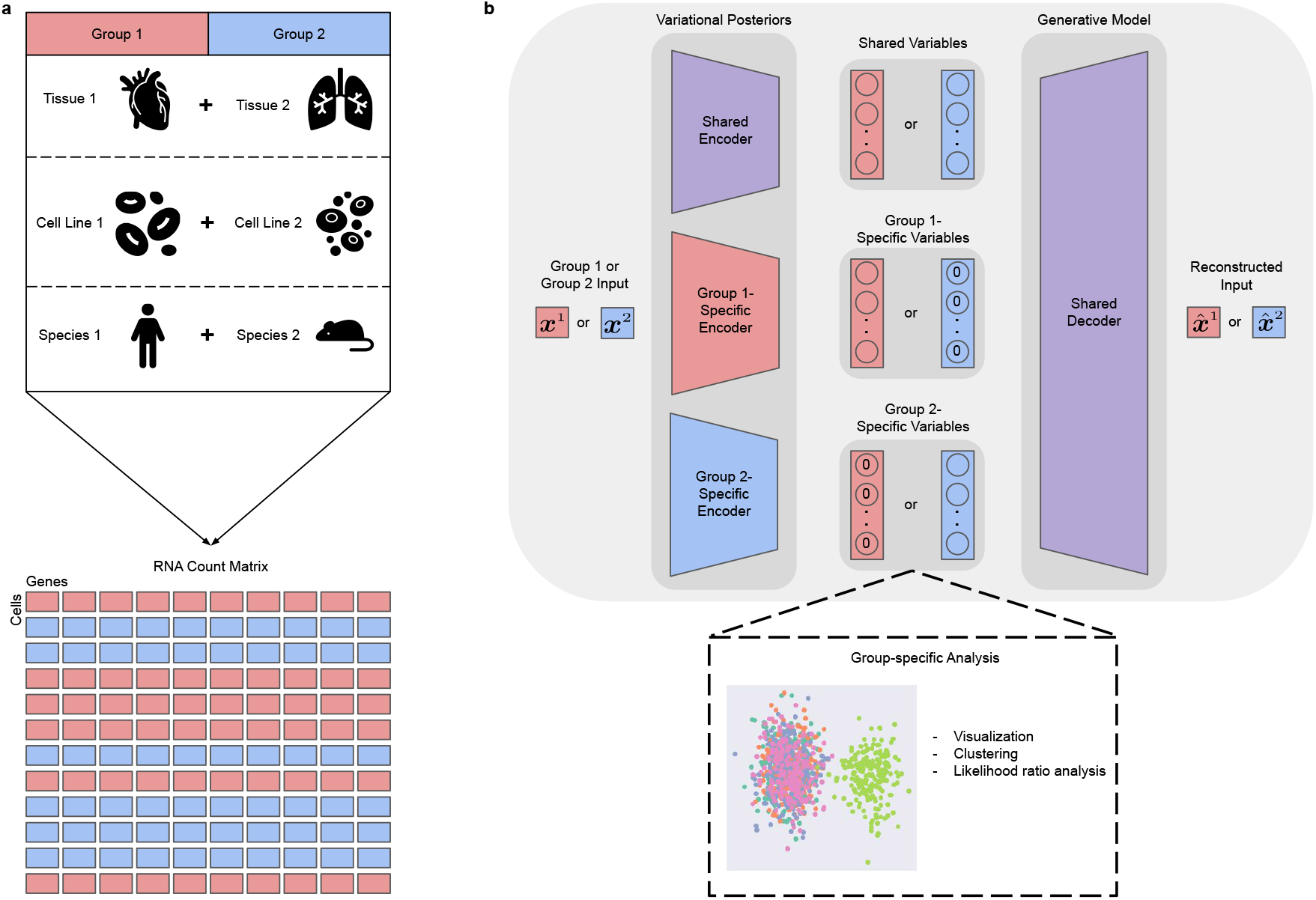
Overview of multiGroupVI. **a**, Given cells divided into non-overlapping groups of interest, multiGroupVI deconvolves the variations shared across groups versus those specific to individual groups. **b**, Schematic of the multiGroupVI model. A shared encoder network embeds cells, regardless of group membership, into the model’s shared latent space, which captures variations shared across all groups. Group-specific encoders also embed cells into group-specific latent spaces, which capture variations particular to a given group. For a cell from a given group *γ*, the group-specific latent variables for other groups *γ*^′^ ≠ *γ* are fixed to be zero vectors. Cells’ latent representations are decoded back to the full gene expression space using a shared decoder. Here for simplicity we depict only two groups, though our model can easily be extended to handle more groups by adding additional group-specific encoders.

The remainder of this paper is organized as follows. First, we formally describe the generative process for multiGroupVI (**Section 2**). We then introduce the inference procedure (**Section 3**), as well as a method for interpreting the group-specific variations captured by multiGroupVI (**Section 4**). We assess the performance of our model in a simulated dataset where ground truth shared and group-specific gene expression patterns are available (**Section 5**). In this setting, our model outperforms state-of-the-art methods for identifying shared versus group-specific variations in the data. Finally, we apply multiGroupVI to re-analyze an existing scRNA-seq dataset [3] (**Section 6**) and recover known cell-type-specific variation of gene expression across tissues.

## 2 The multiGroupVI probabilistic model

We now describe the latent variable model for multiGroupVI. We assume that we have collected gene expression profiles from Γ disjoint groups. That is, for each group *γ* ∈ [Γ], we have *n*_*γ*_ i.i.d. samples 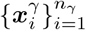 each with measured expression values for *G* genes. We assume that these samples are generated by a random process involving two sets of lower-dimensional latent variables ***z***_*i*_ and 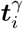 drawn from known prior distributions. Here we denote by ***z***_*i*_ the *shared* latent variables, and we assume that they capture factors of variation shared acrosss all groups. On the other hand, the second set of *group-specific* variables 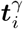 captures factors of variation unique to group *γ*. We place isotropic Gaussian priors on ***z*** and ***t***^*γ*^, i.e.,

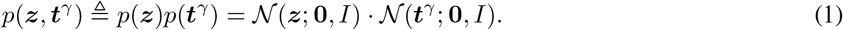

For a given sample 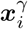 we assume that the group-specific variables for *other* groups (i.e., 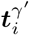 for *γ*^′^≠ *γ*) do not contribute to the generative process and are fixed at 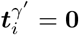 to represent their absence. For notational convenience we define

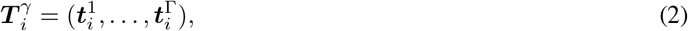

to be the concatenation of all the group-specific variables, including those fixed at **0**. Next, let *f*^*η*^ denote a neural network parameterized by *η* with a softmax output that takes in as input the shared and all of the group-specific variables.

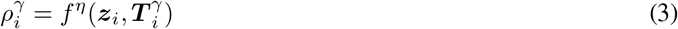

is a vector on the probability simplex that can be viewed as the normalized expression of each gene *g* for the cell 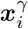. Next, let *ℓ*_*μ*_ and 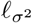 denote the empirical mean and variance of the log-library size. The latent variable

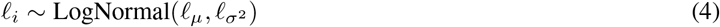

is a scaling factor for each cell that corresponds to discrepancies in sequencing depth and capture efficiency. For each gene *g* ∈ *G*, we assume that the gene expression level 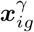 is independent conditioned on 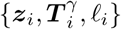 and sampled from an overdispersed count distribution

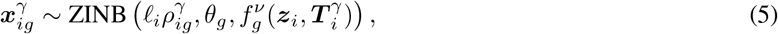

where *θ* ∈ ℝ^|*G*|^ denotes a learned vector of gene-specific inverse-dispersion parameters and *f*^*ν*^ denotes a neural network with parameters *ν* that computes the probability of a gene having zero expression due to dropout effects. We note that our generative process is similar to that of scVI [8], with the addition of the group-specific latent variables. We also note that CPVAE [10] follows a similar, but more restricted, generative process. We expand upon the differences between our generative process and that of CPVAE in **Appendix A**.

## 3 Regularized variational inference

We cannot compute the multiGroupVI posterior distribution directly using Bayes’ rule because the integrals required to compute the model evidence 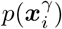 are intractable. We thus instead approximate it with variational inference [18]. More precisely, for cells in a given group *γ*, we approximate the posterior with a factored Gaussian distribution:

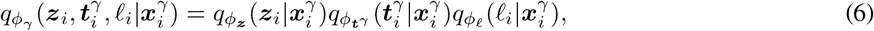

where *ϕ*_*γ*_ denotes the parameters of the approximate posterior. Based on our factorization, *ϕ*_*γ*_ is comprised of three separate components 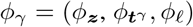. Parameters *ϕ*_***z***_ and *ϕ*_*ℓ*_ are shared across groups in order to encourage 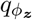 and 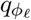 to capture variations shared across all groups and let 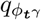 capture variations that are specific to each group *γ*. Following the VAE framework [19], parameters of the approximate posterior for each factor (e.g., mean and variance) are obtained as the output of a deep neural network that takes as input gene expression levels.

We optimize the parameters of the generative model Θ = {*η, ν, θ*}, as well as the ones for the approximate posterior by maximizing the evidence lower bound

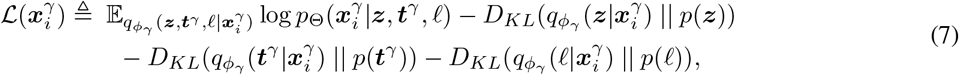

via the stochastic backpropagation algorithm [20]. For our experiments, we use the Pytorch [21] implementation of the Adam optimizer [22] with default hyperparameters.

The group-specific weights 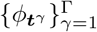 encourage the group-specific latent variables ***t***^*γ*^ to capture variations unique to that group. However, the loss function of (7) does not explicitly prevent ***t*** from also capturing variations shared across groups. Indeed, we find in our experiments on simulated data (**Section 5**) that the group-specific latent variables often capture features shared across populations when learned using (7). To alleviate this issue, we thus need additional constraints. In particular, we extend our prior from (1) to incorporate the group-specific latent variables ***t***^*γ*^*′* for other groups *γ*^′^≠ *γ*, i.e.,

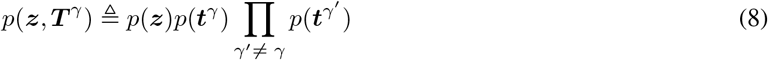

We can then derive a new variational lower bound

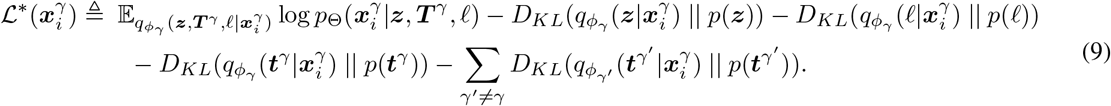

As specified in our generative model, ***t***^*γ*^*′* is fixed at **0** for *γ*^′^ ≠ *γ*. Formally, the corresponding prior *p*(***t***^*γ*^*′*) is a Dirac distribution centered at **0**. However, in that case, the Kullback–Leibler (KL) divergence on the right hand side of (9) is not properly defined (a Gaussian distribution does not admit a density with respect to a counting measure). To work around this issue, we choose to instead use the squared Wasserstein distance [17] as a surrogate to encourage distributional similarity in place of the degenerate KL divergence terms. The Wasserstein distance between a Gaussian random variable and a Dirac distribution has a closed form solution

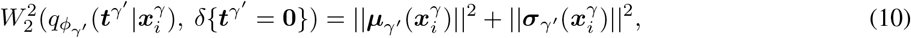

where 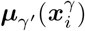 and 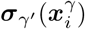 denote the mean and standard deviations of our approximate posterior 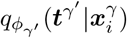. Substituting this expression in for the degenerate KL terms in (9) yields our final objective function

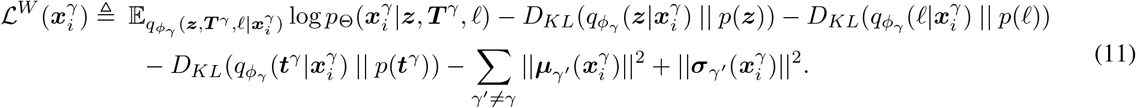

We note that Davison et al. [10] proposed a different regularization scheme to alleviate similar issues found with their CPVAE model. In particular, Davison et al. [10] proposed adding an additional penalty term to their objective function minimizing an upper bound on the mutual information between 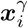 and ***t***^*γ*^*′* for all *γ*^′^ ≠ *γ*. However, in our experiments (**Section 5**) we found their scheme failed to prevent our group-specific variables from capturing undesirable shared variations. We discuss the regularization proposed by Davison et al. [10] in more detail in **Appendix B**.

## 4 Interpreting the group-specific variations captured by multiGroupVI

More complex nonlinear models such as multiGroupVI promise to recover shared and group-specific factors of variation more faithfully than simpler linear models. However, this flexibility comes at the cost of interpretability. While linear models explicitly allow for interpreting the trends captured by the model by manual inspection of its coefficients, (e.g., [23; 24; 13; 7]), such an analysis is not possible for more complex models with neural networks. Thus, in order for practitioners to gain biological insights from multiGroupVI, additional tools are needed.

In this work we consider the task of detecting genes with large group-specific effects, as opposed to those which are primarily governed by the shared latent factors. That is, for cells in a given group *γ* (i.e.,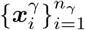), we seek to understand which genes have variations that are captured by the group-specific variables ***t***^*γ*^ as opposed to those that are governed solely by the shared latent variables ***z***.

To do so, we propose the following procedure. Let 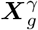 denote the vector of counts for gene *g* across all cells in group *γ*. First, for each cell in a group *γ* we compute the approximate posterior means of latent variables ***z*** and ***t***^*γ*^ using the encoder networks. Next, let ℳ_0_ denote our learned generative model when only given access to the shared latent variables ***z*** (i.e., ***t***^*γ*^ = 0), and let ℳ_1_ denote the model with access to both sets of latent variables (***z, t***^*γ*^). We then compute the logarithm of the likelihood ratio for ℳ_0_ and ℳ_1_ to obtain a Bayes factor

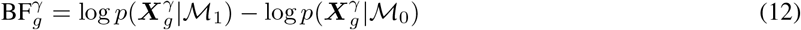

and we rank genes by their Bayes factors, where a larger Bayes factor value indicates greater group-specific effects.

## 5 Performance benchmark on simulated data

We first validated our model using a simulated dataset with known ground truths generated with Splatter [25]. Our dataset consisted of five simulated cell types (**Figure 2a**). Each cell, regardless of cell type, was assigned to one of two groups (**Figure 2b**). Within each group, each cell expressed one of two group-specific gene programs separate from those that distinguished cell types (**Figure 2c**). Thus, each cell’s gene expression profile is composed of shared variations (i.e., those that distinguish cell types) and group-specific variations (i.e., the group-specific gene programs). Our goal is to learn a representation that separates cell types (with strong mixing across groups) in the shared latent space while separating the group-specific gene programs (with strong mixing across cell types) in the group-specific latent spaces. We report hyperparameter values for multiGroupVI and baseline models in **Appendix C** as well as further details describing how this dataset was generated in **Appendix D**. We note that this dataset constitutes a relatively simple multi-group scenario in which each gene feature either exclusively exhibits shared variations or exclusively exhibits group-specific variations. However, we find that, even in this straightforward case, previously proposed methods for disentangling shared and group-specific variations in scRNA-seq data struggle.

**Figure 2:**
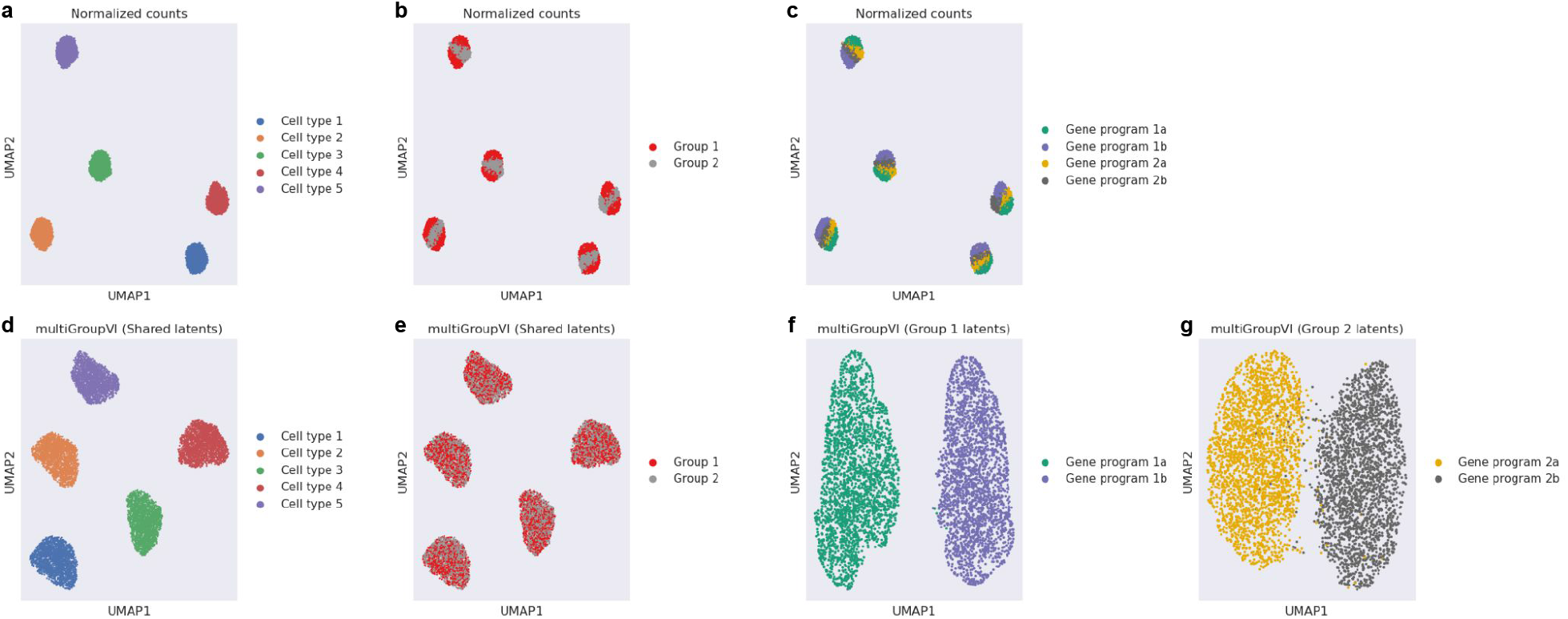
multiGroupVI correctly deconvolves shared and group-specific variations in a simulated dataset. **a-c**, Splatter [25] was used to generate a simulated scRNA-seq dataset. The dataset consisted of cells from five simulated cell types (**a**) divided into two groups (**b**) with each group having two group-specific gene programs (**c**). **d-g**, We find qualitatively that multiGroupVI correctly relegates cell-type-related variations to its shared latent space (**d**) with strong mixing across groups (**e**) while isolating group-specific variations in the group-specific latent spaces (**f, g**).

### 5.1 Evaluation of disentanglement performance

Qualitatively, we find that multiGroupVI separates cells by cell type in its shared latent space (**Figure 2d**) with strong mixing across groups (**Figure 2e**), indicating that the model indeed has captured variations shared across the two groups in its shared latent space. Moreover, we find that multiGroupVI correctly separates the group-specific gene programs in its group-specific latent spaces (**Figure 2f-g**). Taken together, these results demonstrate that multiGroupVI is able to successfully deconvolve shared and group-specific variations.

We benchmarked multiGroupVI’s performance to that of previously proposed methods for analyzing scRNA-seq data (**Figure 3)**. First, to demonstrate the benefits of our nonlinear neural-network-based approach, we compared against a linear model similar to the previously proposed MSFA and GMDF models that we call multi-group factor analysis (MGFA). To demonstrate that similar analyses cannot be performed with trivial combinations of previously existing tools, we also compared against a combination of scVI and MGFA (scVI+MGFA). That is, an scVI model was first trained to capture nonlinear relationships in the data, and then MGFA was run on scVI’s latent representations. We also compared to CPVAE [10] to illustrate the benefits of our more flexible architecture. Finally, to understand the impact of the additional Wasserstein-distance-based regularization term in (11), we trained multiGroupVI models without the additional term (MGVI no reg.) and with the term (MGVI).

**Figure 3:**
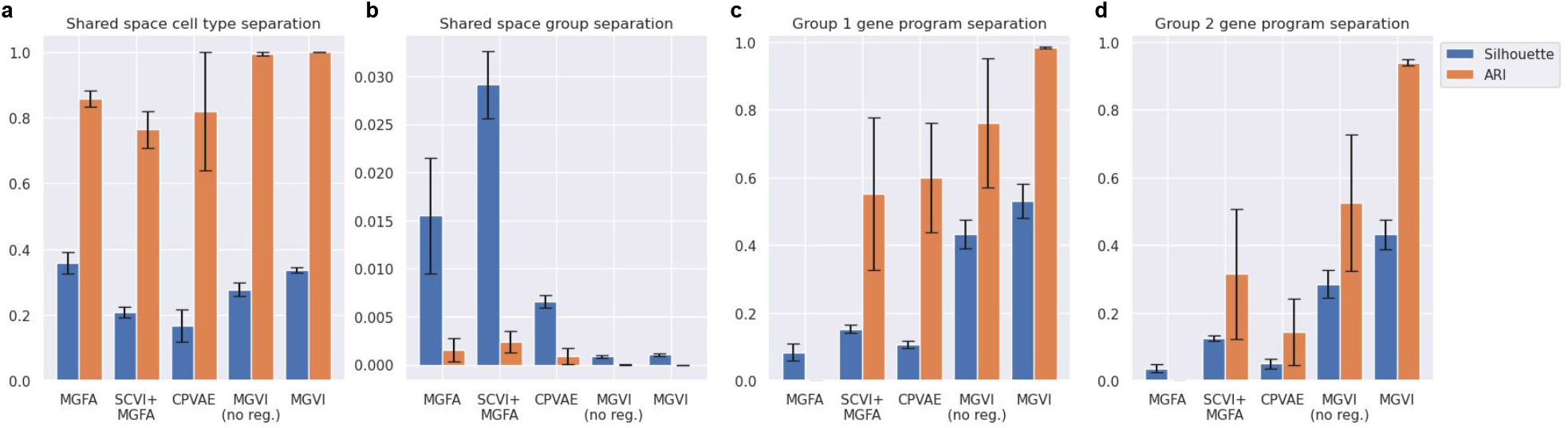
Quantitative benchmarking of multiGroupVI with previous state-of-the-art methods. **a-b**, Separation by cell type (**a**) and group (**b**) in models’ shared latent spaces were quantified using the average silhouette width (Silhouette) and adjusted Rand index (ARI). As cell-type-defining variations are shared across groups, higher values indicate better performance for cell type separation, while lower values indicate better performance for group separation. **c-d**, Separation of cells by group-specific gene programs in models’ group-specific latent spaces was also quantified using the same metrics. Higher values indicate better performance.

For our benchmarking we first quantified how well each method separated cell types (**Figure 3a**) and groups (**Figure 3b**) in their shared latent spaces using average silhouette width (Silhouette) and adjusted rand index (ARI) computed using clusters obtained via *k*-means clustering. For each metric we report the mean and standard error across five random initializations. Since we seek strong separation across cell types and strong mixing across groups, higher values of our metrics indicate better performance for cell type separation while lower values indicate better performance for group mixing. We find that multiGroupVI better separates cell types than previously proposed methods.

We next quantified how well each method separated the group-specific gene programs in their group-specific latent spaces (**Figure 3c-d**). Here we find that only multiGroupVI with the additional regularization term is able to consistently separate the group-specific gene programs in its group-specific latent spaces. On the other hand, the linear MGFA model completely fails to separate the group-specific programs, while the scVI + MGFA workflow, CPVAE, and multiGroupVI with no regularization are far less consistent with their results. Moreover, we found that CPVAE struggled on this task whether trained with its original objective function [10] or our newly proposed regularization term (**Appendix B**), demonstrating the advantages of our more expressive model. Taken together, these results indicate that previously proposed methods for analyzing grouped scRNA-seq are not sufficiently expressive to deconvolve shared and group-specific variations, even in relatively simple multi-group scenarios. On the other hand, we find that multiGroupVI trained with proper regularization successfully disentangled these different sources of variation.

### 5.2 Evaluation of the interpretability procedure

We additionally used our simulated dataset to validate the interpretability procedure proposed in **Section 4**. When generating the dataset, we ensured that genes with cell-type-distinguishing variations shared across groups were completely disjoint from those with group-specific gene program variations. Given that our model appears to correctly isolate group-specific variations in its group-specific latent spaces (**Figure 2)**, we would thus expect genes with the highest likelihood ratios to be those genes expressing group-specific variations.

We report our results in **Figure 4**. We find that the top genes as ranked by our likelihood ratio ranking indeed are those known to express group-specific variations. We compared the performance of our likelihood ratio ranking with two other methods designed for interpreting the latent spaces of unsupervised machine learning models: Hotspot [28], a method that ranks genes by spatial autocorrelation when provided a given metric of cell-cell similarity (e.g. the latent space of an autoencoder), and an adaptation of integrated gradients [26] for unsupervised models proposed in Crabbé and van der Schaar [27] applied to multiGroupVI’s group-specific latent spaces. We find that neither of these baseline methods is as effective at recovering genes with group-specific variations as our likelihood ratio ranking.

**Figure 4:**
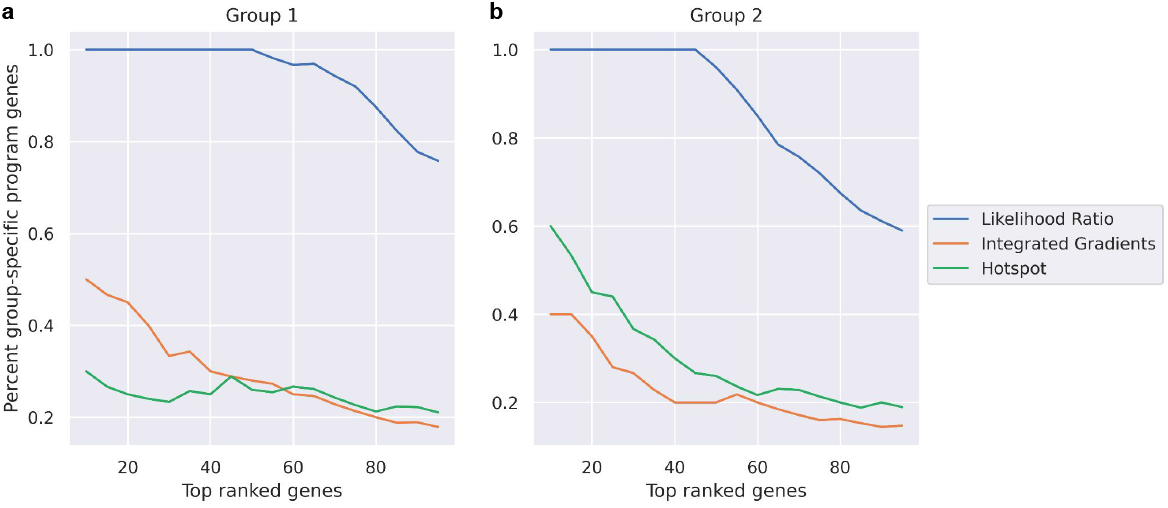
Benchmarking methods for recovering genes with largest group-specific effects in multiGroupVI. **a-b**, Multiple methods (the proposed likelihood ratio ranking, integrated gradients [26; 27], and Hotspot [28]) were applied to rank the genes in our simulated dataset by how strongly they were captured by multiGroupVI’s group-specific latent spaces. Here we consider the top *n* genes ranked by each method for varying values of *n*, and we plot the percentage of returned genes known *a priori* to have group-specific variations for our simulated groups 1 (**a**) and 2 (**b**). Higher values indicate better performance. We find that our likelihood ratio ranking recovers far more genes with known group-specific effects than other methods.

## 6 Application to real-world data

We next applied multiGroupVI to explore a real-world scRNA-seq dataset from Haber et al. [3]. In particular, we consider a dataset consisting of 11, 665 epithelial cells from three regions of the mouse small intestine: the duodenum, jejunum, and ileum. While all the regions of the small intestine contribute to the same high-level process (i.e., the absorption of nutrients), each region is also known to have specific functions. For example, it is well known [29; 30] that vitamin B_12_ is absorbed primarily in the ileum. To better understand these region-specific processes, it is thus of interest to separate variations in gene expression shared across regions versus those specific to individual regions. In our experiments here we trained multiGroupVI using the region of origin as the group label for each cell, and we present our results in **Figure 5**. We provide details on our preprocessing of this dataset in **Appendix E**.

**Figure 5:**
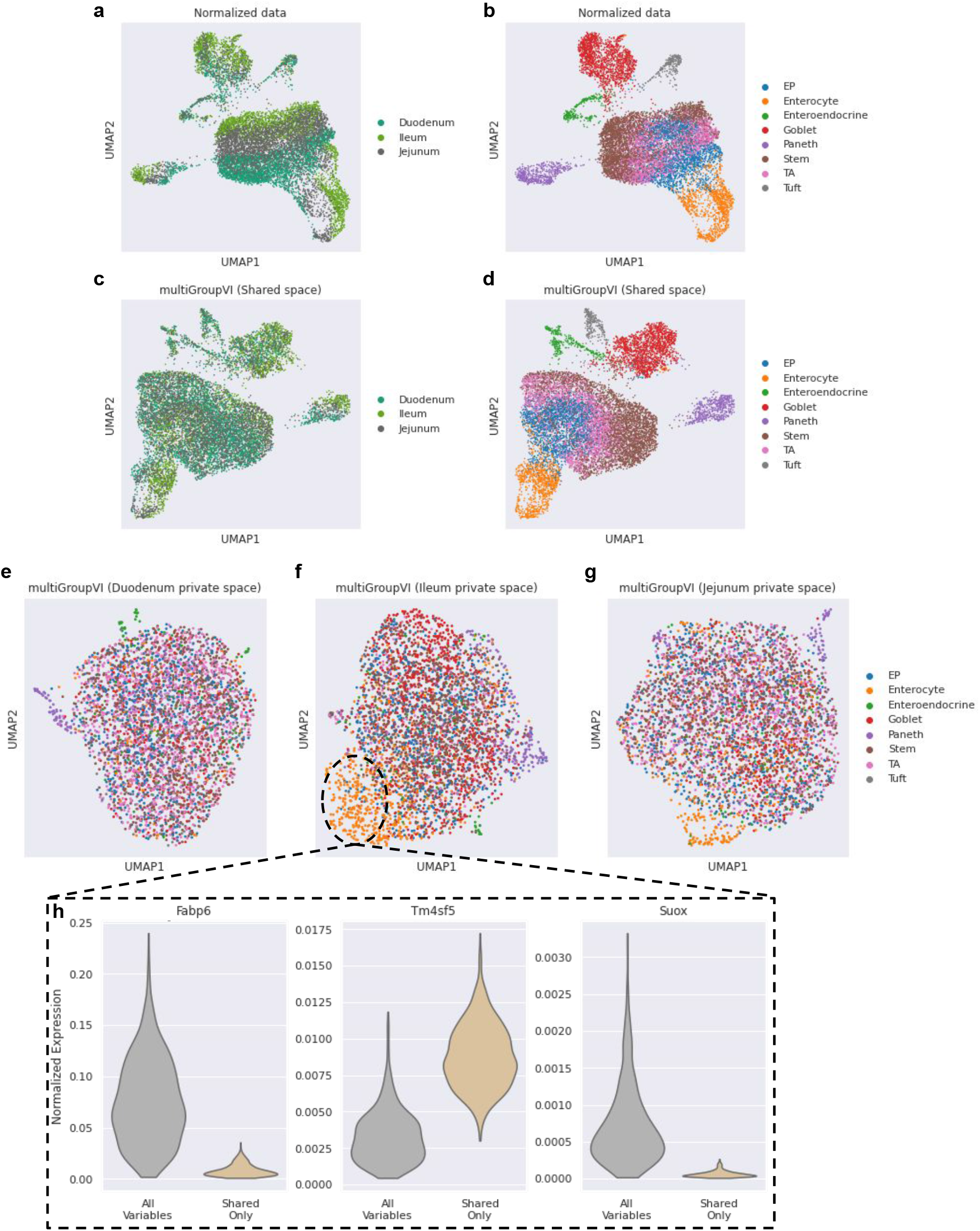
Exploring the multi-region mice intestine epithelial cell dataset from Haber et al. [3] with multiGroupVI. **a-b**, The original data (i.e., normalized and log-transformed counts) strongly separated by both region (**a**) and cell type (**b**). **c-d**, In the multiGroupVI shared latent space we observe stronger mixing across regions (**c**) while separation between cell types is preserved (**d**). **e-g**, In the region-specific multiGroupVI latent spaces we observe some separation between cell types, indicating that region-specific variations are dependent on cell type. For example, we find that enterocytes strongly separate from other cell types in the ileum-specific latent space. **h**, Distributions of normalized expression values 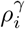 for ileum enterocytes as computed by the multiGroupVI decoder when decoding using all latent variables versus only the shared variables for the top three genes ranked by likelihood ratio for ileum enterocytes.

While cells originally strongly separated based on both region (**Figure 5a**) and cell type (**Figure 5b**), we observe mixing across regions in the multiGroupVI shared latent space (**Figure 5c**) with cells separating primarily based on cell type (**Figure 5d**). We next inspected the region-specific multiGroupVI latent spaces (**Figure 5e-g**). We find that cell types tend to mix in the region-specific latent spaces, with some notable exceptions. For example, we find that Paneth cells separate from other cell types in all of the region-specific latent spaces. This separation indicates the presence of Paneth-cell-specific variations unique to each region, which was also noted by [3]. Similarly, in the ileum and jejunum-specific latent spaces (**Figure 5f**), we find that enterocytes strongly separate from other cell types, indicating the presence of enterocyte-specific variations found only in those regions.

Using ileum enterocytes as a case study, we next applied the likelihood ratio procedure described in **Section 4** to explore the underlying biology driving their separation from other cell types in the ileum-specific latent space. After computing gene rankings, we plotted the distributions of normalized expression values as computed by the multiGroupVI decoder (i.e., the entries of 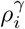 from (3)) when using all latent variables versus only the shared latent variables (**Figure 5h**) for the three most highly ranked genes. The significant differences in the distributions of these values suggest that these genes exhibit strong ileum-specific expression patterns in enterocytes. On the other hand, we find that normalized expression values for enterocytes in the other regions do not significantly differ whether computed using all latent variables or using only the shared variables (**Figure 1)**. This result suggests that, for enterocytes in other regions, these genes’ expression levels are solely governed by the shared latent factors ***z*** without additional region-specific effects. Taken together, our results indicate that these genes likely are involved in ileum-specific enterocyte functions.

We then compared the top genes returned by our procedure to those originally found by Haber et al. [3] to have ileum-specific variations in enterocytes. We find that our results (**Table 1)** largely agree with those of [3], with most of the top genes as ranked by our likelihood ratio procedure also found by the original study to have significant region-specific differences. We further inspected (**Figure 2)** the genes that were highly ranked by multiGroupVI but not originally highlighted in [3], and found that these genes indeed appear to exhibit region-specific variations in enterocytes. This indicates that multiGroupVI potentially captured region-specific enterocyte variations missed in the original study. Altogether, these results demonstrate that multiGroupVI can be used both to identify the presence of group-specific gene expression patterns in scRNA-seq data, as well as to identify the specific genes underlying these variations.

## 7 Discussion

In this work we introduced multiGroupVI, a deep generative model designed to isolate latent factors of variation shared across cells from different pre-defined groups from those that are specific to individual groups. We first validated our approach on a simulated dataset with known ground truths, and we found that it successfully separated shared and group-specific variations while less expressive models failed to do so. We also applied multiGroupVI to analyze an scRNA-seq dataset consisting of epithelial cells from different regions of the mouse small intestine and found that it could identify cell-type-specific variations unique to individual regions.

Our initial results with multiGroupVI presented in this manuscript suggest that it indeed can deconvolve shared and group-specific variations more effectively than previously proposed methods. However, interpreting the variations captured by the different multiGroupVI latent spaces remains a challenge. In this work, we presented a procedure for identifying genes with strong group-specific effects. While we found this procedure to be effective, it is limited to identifying single genes as opposed to coordinated gene programs that may be more biologically meaningful. Moreover, our procedure is limited to identifying genes with group-specific effects and does not provide further insights on the trends captured in the model’s shared latent space. In practice, these shared variations may be of more interest than the group-specific effects, such as in previous studies investigating pan-cancer gene programs [7; 4]. For future work we thus plan on augmenting our multi-group architecture with recent advances in interpretable VAE models, such as BasisVAE [31], which identifies clusters of coordinated features captured by the learned latent variables in addition to performing dimensionality reduction.

Beyond the datasets considered in this work, multiGroupVI can easily be applied to other single-cell data analysis problems in which shared versus group-specific disentanglement is desired. For example, when given expression levels of orthologous genes from multiple species, multiGroupVI could disentangle variations shared across species versus those that are specific to an individual species. Furthermore, when analyzing multi-omic datasets, it may be of interest to decompose the data into sets of latent variables that reflect variations shared across modalities versus those that are specific to a single modality. Previous work has addressed this problem with linear models [32]; however, to our knowledge no works have applied more expressive deep learning modeling techniques to this problem. Indeed, previously proposed deep learning models for multimodal single-cell data, such as totalVI [33] and MultiVI [34], learn only a single set of latent variables shared between all modalities and are not designed to deconvolve shared versus modality-specific variations. For future work we will thus explore extending these models to incorporate modality-specific latent variables inferred via modality-specific encoders trained with a disentanglement-promoting loss function similar to (11).

Moreover, beyond the basic shared versus group-specific structure considered in this work, our model could easily be extended to handle richer sets of latent variables reflecting additional assumptions on the underlying structure of the data. For instance, multiGroupVI could be extended to include pairwise sets of latent variables that capture variations shared in only a subset of groups. Finally, future work will explore using more efficient encoding architectures for learning different sets of latent variables. For example, rather than using an entirely separate encoder network for each group of latent variables, it may be sufficient to use a single encoder network with shared initial layers followed by group-specific heads.

## Acknowledgements and Disclosure of Funding

We thank Tara Chari for feedback throughout the duration of this project that greatly improved this work. We also thank members of the Regev Lab and the Artificial Intelligence/Machine Learning department at Genentech for providing constructive feedback on earlier versions of the results presented in this work. We warmly thank Su-In Lee and Chris Lin for valuable discussions on disentanglement for deep generative models as well as interpretability for machine learning models for biological data. We also acknowledge the MLCB reviewers whose feedback helped improve this work. Finally, we acknowledge the developers/maintainers of software packages in the scverse consortium^1^ - including scanpy [35] and scvi-tools [1] - for their development of high-quality open-source software packages for analyzing single-cell omics data which were used in this work. This material is based upon work supported by the National Science Foundation Graduate Research Fellowship under Grant No. DGE-214000.

## Code Availability Statement

We implemented multiGroupVI in PyTorch [36] with the scvi-tools [1] library using the base model in the scvi-tools-skeleton repository^2^ as a starting point. Some code relating to data loaders for training multiGroupVI models was adapted from code from the official implementation of the contrastiveVI [14] model^3^. Our implementation is available at https://github.com/Genentech/multiGroupVI and is released under the Apache 2.0 license.

## Appendices

The appendices are organized as follows. In **Appendix A** we provide additional details on CPVAE generative process from Davison et al. [10]. In **Appendix B** we discuss the differences between our proposed Wasserstein-distance-based regularization and the regularization proposed in Davison et al. [10] to penalize the leakage of information between shared and group-specific latent spaces. In **Appendix C** we provide information on the hyperparameters used for multiGroupVI and baseline models in our experiments. In **Appendix D** and **Appendix E** we provide further details on the generation of our simulated dataset and the preprocessing of our real-world dataset, respectively. Finally, in **Appendix F** and **Appendix G** we provide the supplementary figures and tables, respectively, that are referenced in the main text.

### A Further details on the CPVAE [10] generative process

In a previous work, Davison et al. [10] proposed the cross-population variational autoencoder (CPVAE) model with a similar goal of isolating variations shared across groups versus those specific to individual groups. Here we present CPVAE in more detail, and highlight notable differences between CPVAE and our proposed multiGroupVI model. As with multiGroupVI, the CPVAE generative process assumes that each sample is generated from two sets of latent variables: shared factors ***z*** as well as group-specific factors ***t***^*γ*^. CPVAE also places isotropic Gaussian priors on these variables, i.e.,

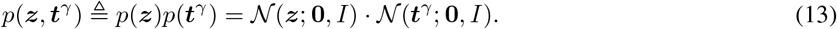

Conditioned on the latent variables, samples are generated from a distribution

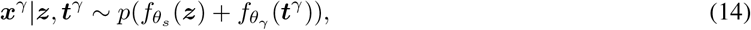

where *p* is an appropriately chosen distribution for modeling the observed data, 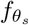 is a nonlinear function with parameters *θ*_*s*_ shared across groups that outputs a vector of parameters for *p*, and 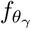 is a similar nonlinear function with group-specific parameters *θ*_*γ*_. We highlight that this modeling choice is much more restrictive compared to multiGroupVI: while CPVAE forces the parameters of *p* to be a simple sum of arguments, multiGroupVI allows for the parameters of *p* to be arbitrary functions of ***z*** and ***t***^*γ*^.

In Davison et al. [10], the authors choose *p* to be a Gaussian distribution. However, previous work [11; 12] has demonstrated that using a distribution that more faithfully models the technical biases and noise characteristics of scRNA-seq data is crucial for learning effective representations of the data. Thus, in our experiments applying CPVAE to scRNA-seq data, we choose *p* to be the ZINB distribution.

Specifically, as with multiGroupVI we first assume the presence of an additional latent variable

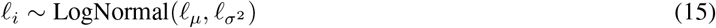

representing a scaling factor for each cell. Next, we let 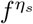 denote a neural network with parameters *η*_*s*_ shared across groups with a softmax output, and we let 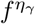 be defined analogously but with group-specific parameters *η*_*γ*_.

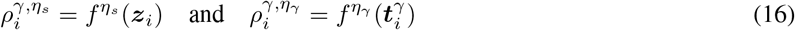

are vectors on the probability simplex that can be viewed as the normalized expression of each gene *g* for the cell 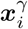.

For each gene *g* ∈ *G*, we assume that the gene expression level 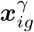 is independent conditioned on 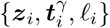 and sampled from

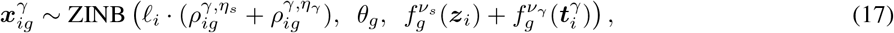

where *θ* ∈ ℝ^|*G*|^ denotes a learned vector of gene-specific inverse-dispersion parameters and 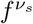 and 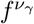 denote neural networks with parameters *ν*_*s*_ and *ν*_*γ*_ that compute the probability of a gene having zero expression due to dropout effects.

### B Further details on the CPVAE regularization proposed in Davison et al. [10]

As was also found in our experiments with multiGroupVI, Davison et al. [10] found that information tended to “leak” between latent spaces (i.e., shared variations were captured in group-specific latent spaces and vice-versa) without additional constraints on their model. To mitigate this problem, the authors proposed adding an upper bound on the sums of the mutual information between ***x***^*γ*^ and ***t***^*γ*^*′* for groups *γ*^′^ ≠ *γ* as a penalty to their ELBO function. In particular the authors derive the following bound

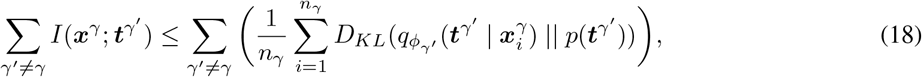

and take *p*(***t***^*γ*^*′*) = 𝒩 (***t***^*γ*^*′* ; **0**, *I*). That is, for samples from a group *γ*, the regularization of Davison et al. [10] encourages the distribution of group-specific latent variables for other groups *γ*^′^ to be “uninformative” zero-mean, unit-variance Gaussians. We note this choice of an “uninformative” prior is far more permissive than our more strict choice of a Dirac delta distribution centered at **0**.

The authors initially found that adding this term naively to their ELBO did not prevent leakage between the latent spaces. To remedy this, the authors then proposed to eliminate the KL divergence term 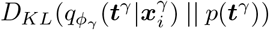 from their objective function. That is, for samples from group *γ*, the final objective function of [10] no longer encourages the distribution of the true group-specific variables to be close to an isotropic Gaussian prior. The authors justified this choice by stating that the additional flexibilty given to the true group-specific variables ***t***^*γ*^ (as compared to ***t***^*γ*^*′* for *γ*^′^ ≠ *γ*) would enable them to better capture the true group-specific variations. However, removing this term is not theoretically justified.^4^ Moreover, it effectively results in no prior distribution being placed on ***t***^*γ*^, which may be problematic for downstream applications of the model. Thus, for our experiments involving the regularization proposed in [10], we use the penalty term of (18) while leaving the KL divergence term 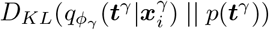 intact.

We found that this regularization with the intact KL term indeed failed to mitigate the leakage problem. However, we notably also found that our proposed Wasserstein regularization also failed to improve CPVAE’s performance. In **Figure 3** we produce the benchmarking metrics presented in **Figure 3** for CPVAE trained with different regularizations: no regularization (CPVAE No reg.), the regularization of Davison et al. [10] (CPVAE Davison) and with our Wasserstein distance penalty (CPVAE Wass.).

To better understand whether this lack of improvement was due to the less flexible CPVAE generative process or issues with the regularization terms, we also (**Figure 4)** trained multiGroupVI models with the same regularization penalties, i.e.: no regularization (MGVI No reg.), the regularization proposed in Davison et al. [10] (MGVI Davison), and our proposed penalty (MGVI). We find that our Wasserstein regularization significantly improved the performance of multiGroupVI, while the Davison et al. [10] regularization failed to do so. Taken together, these results illustrate both the necessity of our more flexible architecture as well as the advantages of our regularization approach over that of [10].

### C Hyperparameter choices for multiGroupVI and baseline models

For both the simulated and real-world datasets multiGroupVI and baseline models were trained with 10-dimensional shared latent spaces and 10-dimensional group-specific latent spaces. All models were trained for 500 epochs using the PyTorch [21] implementation of the Adam optimizer [22] with default hyperparameters. For deep learning methods (i.e., multiGroupVI and CPVAE), we used the default encoder and decoder architectures from the scvi-tools library. Specifically, encoder networks had one hidden layer with 128 units trained with a dropout rate of 0.1, while decoder networks had the same architecture in reverse.

### D Further details on simulated data generation

To generate our simulated dataset, we used the Splatter [25] model. However, as Splatter was not designed to simulate data generated from multiple distinct sets of latent factors, we were required to generate our dataset in multiple steps.

First, we simulated the expression of 5, 000 genes for 10,000 cells divided into five cell types. When generating expression values for these genes we used the default Splatter parameters, and cells were assigned to cell types with equal probability. These 5,000 simulated genes represent genes with variations that are shared across groups.

Next, we simulated the expression of 200 additional genes for the same 10, 000 cells. For these genes, we chose a smaller value of 6 for the location parameter for the distribution of library sizes instead of the default value of 11; all other parameters were left at their default values. These genes represent more subtle variations that are specific to different groups of cells.

Specifically, when generating expression values for these genes we used Splatter to simulate four “subgroups” of cells, where here subgroups are equivalent to cell type labels returned by Splatter. As with cell types in the initial set of 5, 000 genes, cells were assigned randomly to subgroups with equal probability. The first two of these subgroups were combined to form one larger group (Group 1), with the two individual subgroups representing group-specific gene programs. Similarly, the remaining two subgroups were combined to form another group (Group 2), with the individual subgroups again representing group-specific gene programs.

The expression values of the 5,000 original genes and the 200 additional genes were then concatenated together to form our final dataset.

### E Preprocessing for real-world single-cell data from Haber et al. [3]

The mouse small intestine epithelial cell dataset is publicly available from the NIH Gene Expression Omnibus (GEO) as GSE92332. In particular, the file GSE92332_Regional_UMIcounts.txt.gz available at https://www.ncbi.nlm.nih.gov/geo/query/acc.cgi?acc=GSE92332 contains the raw count matrix for the data considered in this study. The full count matrix was subsetted to 2,000 highly variable genes using the scanpy [35] implementation of the Seurat v3 [37] highly variable gene selection procedure. Cell type and region metadata labels for each cell were provided by the authors of Haber et al. [3]. Jupyter notebooks with our full code for preprocessing the dataset are available at https://github.com/Genentech/multiGroupVI/tree/main/notebooks.

### F Supplementary Figures

**Supplementary Figure 1:**
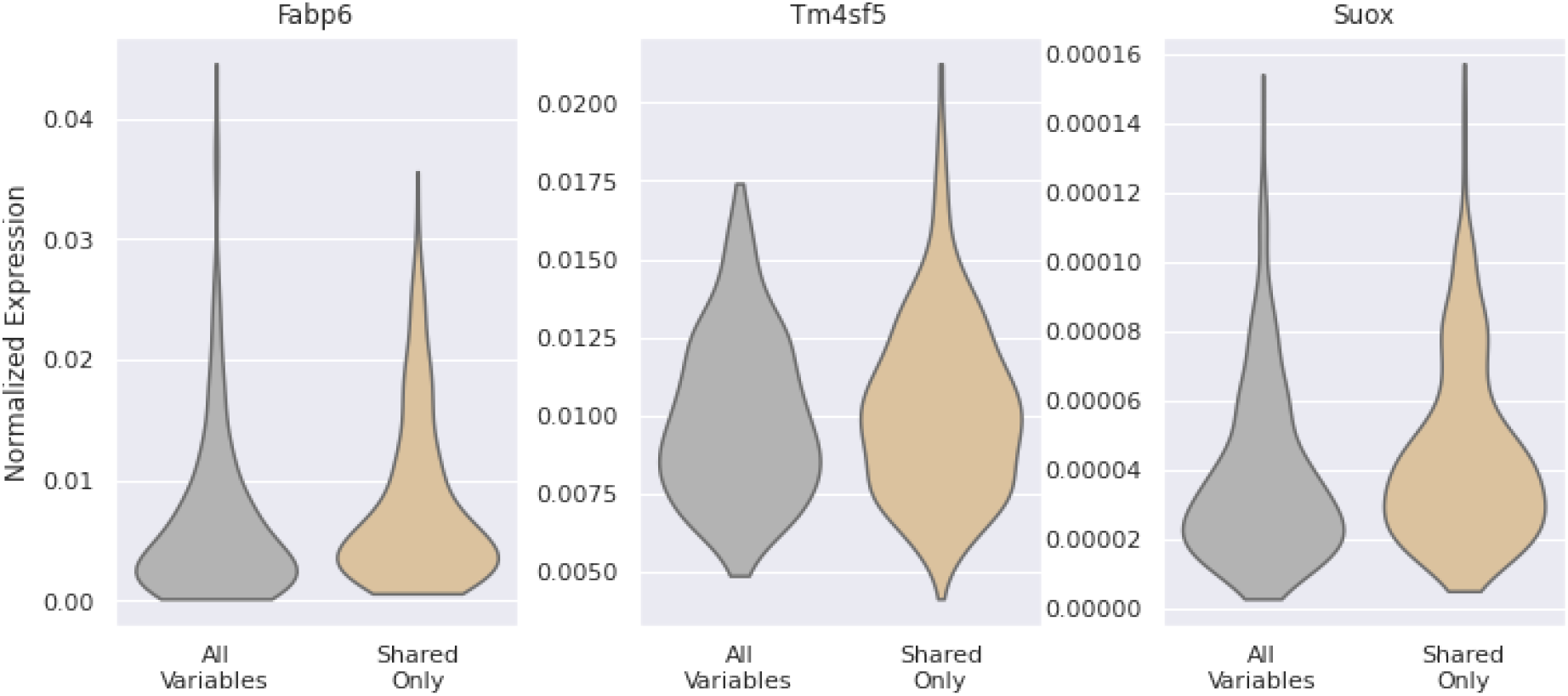
Distributions of multiGroupVI normalized expression values 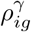 for non-ileum enterocytes of top genes ranked by likelihood ratio for ileum enterocytes. Here we show distributions of the normalized expression values for non-ileum enterocytes computed by the multiGroupVI decoder using all latent variables versus only the shared variables. Genes shown are the top three genes ranked by likelihood ratio for ileum enterocytes. We find for non-ileum enterocytes that the distributions of normalized expression values do not significantly differ when using all variables versus only shared variables.

**Supplementary Figure 2:**
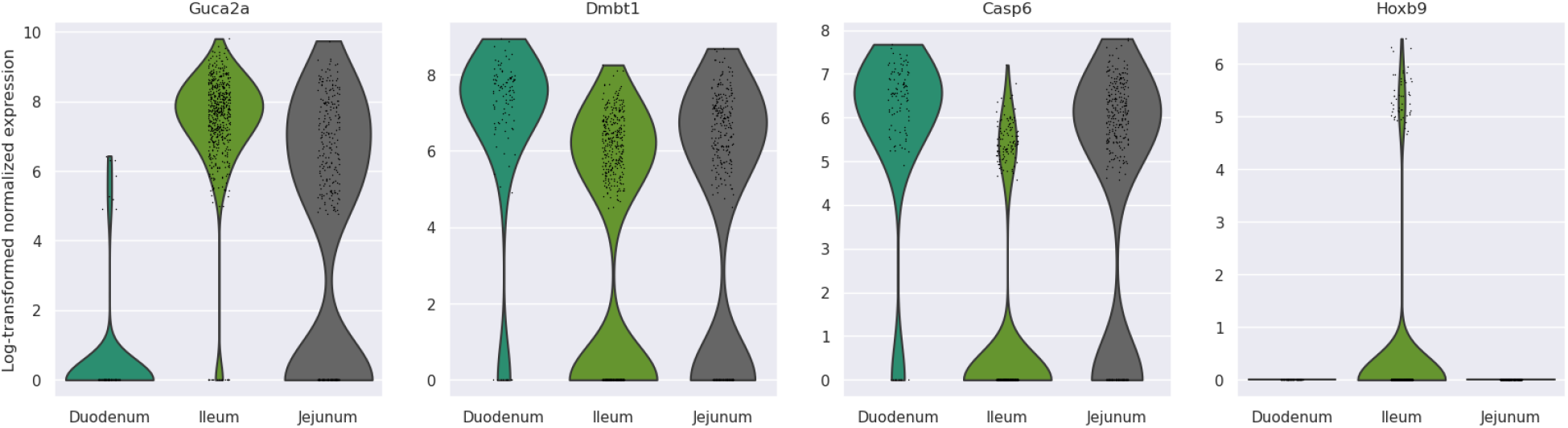
Distributions of expression values in enterocytes for genes found to have ileum-specific enterocyte variations using multiGroupVI but not noted in Haber et al. [3]. Values shown are log-transformed normalized counts computed using a standard scanpy [35] workflow. We find that the distributions of these genes’ expression values in enterocytes do indeed appear to exhibit region-specific trends, despite not being originally noted in [3].

**Supplementary Figure 3:**
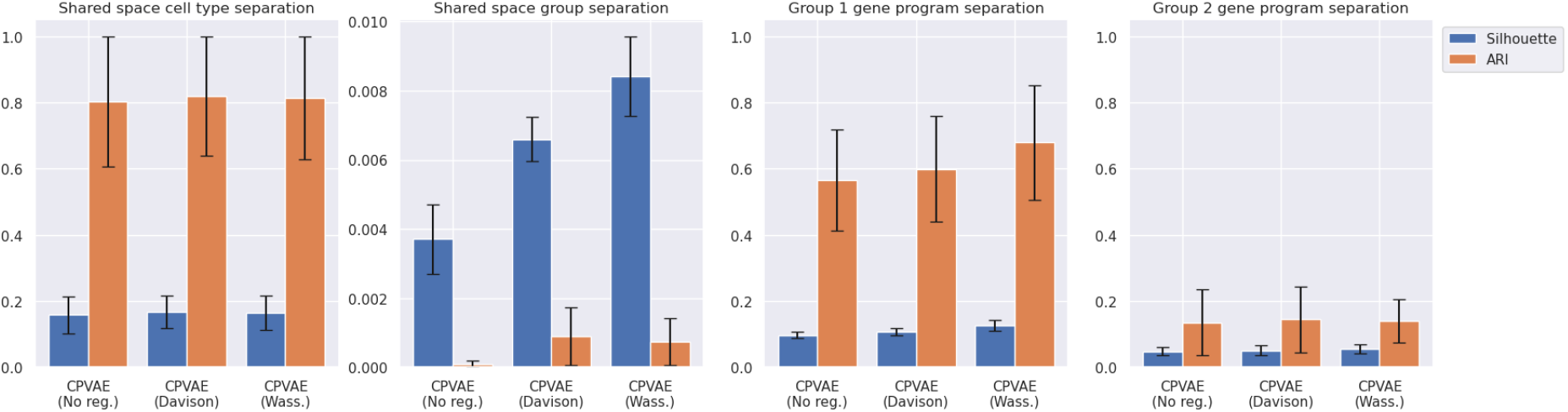
Simulated data benchmark results for CPVAE trained with different regularizations. We find that CPVAE’s performance does not vary significantly on our simulated data benchmarks when trained without regularization (CPVAE No reg.), with the regularization proposed in Davison et al. [10] (CPVAE Davison), or with our proposed Wasserstein distance regularization (CPVAE Wass.).

**Supplementary Figure 4:**
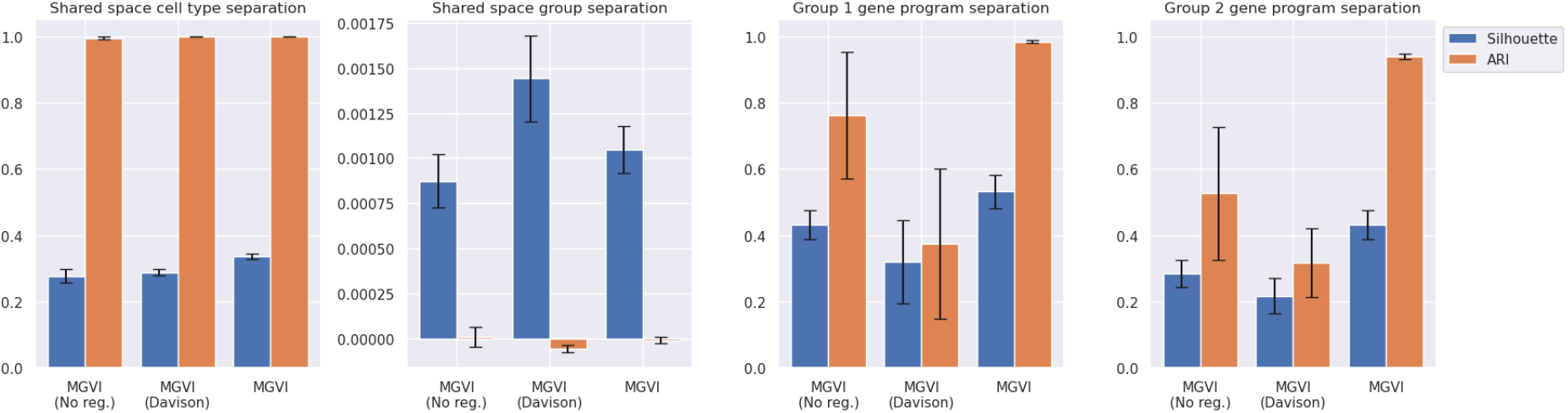
Simulated data benchmark results for multiGroupVI trained with different regularizations. We find that multiGroupVI’s performance is significantly improved by our proposed Wasserstein distance regularization (MGVI), while the regularization proposed by Davison et al. [10] did not improve our model’s performance (MGVI Davison) compared to no regularization (MGVI No reg.).

### G Supplementary Tables

**Supplementary Table 1:** Top 20 genes with ileum-specific enterocyte variations as ranked by our proposed multi-GroupVI interpretability procedure. Genes in green were also found by Haber et al. [3] to have region-specific variations in enterocytes. We find substantial overlap between our list and the findings of Haber et al. [3].

| Gene Name |
| --- |
| Fabp6 |
| Tm4sf5 |
| Suox |
| Guca2a |
| Slc51a |
| Rbp2 |
| Prap1 |
| Mep1a |
| Smim24 |
| Fabp1 |
| Arg2 |
| Tmigd1 |
| Fabp2 |
| Dmbt1 |
| Lgals3 |
| Casp6 |
| Aldob |
| Cda |
| Hoxb9 |
| Khk |

## Footnotes

1 https://scverse.org/

2 https://github.com/scverse/scvi-tools-skeleton

3 https://github.com/suinleelab/contrastiveVI

4 See comments at https://openreview.net/forum?id=r1eWdlBFwS.

